# Salt stress shifts wild tomato root systems toward transport efficiency along a gravitropism-constrained Pareto front

**DOI:** 10.64898/2026.07.16.738932

**Authors:** Aaron Garza, Kathryn Altman, Alleen Koenig, Kendall Richards, Ruth Nichols, Maryam Rahmati-Ishka, Guillaume Lobet, Magdalena M. Julkowska, Arjun Chandrasekhar

**Author notes:** Authors for correspondence: Arjun Chandrasekhar; Magdalena M. Julkowska.

## Abstract

- Root system architecture balances the carbon cost of building roots (wiring cost) against transport speed to the shoot (conduction delay). Wild tomato (*S. pimpinellifolium*) roots lie near the Pareto front, but existing models treat lateral roots as straight, misrepresenting curved architectures.
- We extended the Pareto model to account for lateral-root curvature, estimating curvature parameter *G* and trade-off weight *α* for 1,597 root systems from 240 accessions grown with or without 100 mM NaCl. We also measured root angles and conducted a GWAS. This is the first Pareto model of root architecture to incorporate lateral-root curvature and the first study to demonstrate heritability of position on the Pareto front and identify an associated locus.
- Curvature improved model fit in 80% of root systems after penalizing the additional parameter. Salt shifted root systems toward conduction-delay minimization (lower *α*) across all four architectural types. Lateral roots grew 10° closer to vertical, while main-root angle was unchanged. *α* was more heritable under salt, and a chromosome 6 locus within an intron of the *ISOAMYLASE 3* ortholog explained approximately 11% of its variation.
- Salt reorganizes lateral roots, but not the main root, toward faster transport, with a heritable, mappable component.

## 1 Introduction

Root system architecture determines how a plant explores soil and how efficiently water and nutrients reach the shoot (Lynch, 1995; Rangarajan et al., 2018; Schneider and Lynch, 2020). Because every millimeter of root costs carbon to build and maintain, architecture reflects a compromise between the resources invested in the network and the performance the network delivers. Wild tomato (*S. pimpinellifolium*) is a useful system for studying this compromise: its accessions span coastal and inland habitats in Ecuador and Peru, differ markedly in root architecture, and respond to salt stress with characteristic changes in lateral root number, length and angle (Julkowska et al., 2014; Morton et al., 2024; Rahmati Ishka et al., 2026).

Network optimality theory provides a quantitative way to describe this compromise. Root systems, like neuronal arbors and shoot vascular networks, can be represented as graphs that balance two competing objectives: minimizing *wiring cost*, the total length of material in the network, and minimizing *conduction delay*, the path length from each tip to the base (Conn et al., 2017a,b; Chandrasekhar and Navlakha, 2019). Because both objectives cannot be minimized at once, optimal architectures form a Pareto front, and a single trade-off weight *α* locates any architecture along it. We previously showed that *S. pimpinellifolium* root systems lie close to this front and that four visually scored architectural types (Telephone Pole, Droopy Telephone Pole, Christmas Tree and Broomstick) occupy distinct regions of it (Chandrasekhar and Julkowska, 2022).

That framework, like all previous implementations, assumed that lateral roots connect to the main root along straight lines. Lateral roots do not grow straight. They emerge at a wide angle and then bend toward a species- and condition-specific gravitropic setpoint angle (GSA) (Digby and Firn, 1995), a trajectory controlled by auxin and opposed by cytokinin (Rosquete et al., 2013; Roychoudhry et al., 2013; Waidmann et al., 2019). A curved root is longer than the chord between its ends, so curvature raises both wiring cost and conduction delay. A straight-line model therefore estimates *α* from the wrong set of feasible architectures, and it cannot represent changes in root direction, which are among the most conspicuous responses of roots to salt (Sun et al., 2008; Galvan-Ampudia et al., 2013).

Here we extend the Pareto model so that each lateral root follows a curved trajectory, and we ask three questions. First, does accounting for curvature improve the description of real root systems enough to justify the extra parameter? Second, how does salt stress move *S. pimpinellifolium* root systems within the curvature-constrained trade-off space? Third, is the position of a root system on the front heritable and genetically mappable? We combine the model with direct measurements of root angle from the same tracings, so that each model-based conclusion can be checked against the raw geometry, and with a GWAS across 189 accessions.

## 2 Materials and Methods

### 2.1 Plant material and growth conditions

All experiments were conducted on vertical square agar plates. Seeds were surface-sterilized for 10 min in 50% bleach, rinsed five times with autoclaved milli-Q water, and germinated on [iinline] strength Murashige and Skoog (MS) medium supplemented with 0.5% (w/v) sucrose, 0.1% (w/v) MES and 1% (w/v) agar, pH 5.8 (Rahmati Ishka et al., 2026; Ishka et al., 2026). After 24 h of stratification at 4 °C in the dark, plates were placed vertically in a Conviron growth chamber at 130–150 µmol m^−2^ s^−1^, 20 h light / 4 h dark, 25 °C day / 20 °C night and 60% relative humidity. Four days after germination, seedlings were transferred one per plate to fresh [iinline] MS medium with or without 100 mM NaCl.

The natural diversity panel comprised *S. pimpinellifolium* accessions grown in six experimental batches (PimpiBig1–6) with four biological replicates per accession and condition (Rahmati Ishka et al., 2026). Plates were scanned (EPSON flatbed scanner) every 24 h for five days after transfer. Only *S. pimpinellifolium* accessions were analyzed; cultivated tomato (*S. lycopersicum*) accessions included in the original experiments were excluded. *S. pimpinellifolium* accessions were identified as those listed in the KAUST–TGRC accession table; three accession codes absent from it were excluded.

### 2.2 Tracing root system architecture

The main root and all visible lateral roots were traced manually on the last-day image of each plate using SmartRoot (Lobet et al., 2011), and each root was exported as an ordered series of *x*–*y* coordinates. Each root system was also assigned to one of four architectural types (Telephone Pole, Droopy Telephone Pole, Christmas Tree or Broomstick) by visual inspection (Chandrasekhar and Julkowska, 2022). The analysis set comprised 1,597 last-day root systems from 240 accessions (814 control, 783 salt); 1,385 of these carried one of the four architectural-type labels.

### 2.3 Curvature-constrained Pareto model

We represent a root system as a weighted, connected, acyclic graph *A* = (*V, E*) in which vertices are two-dimensional coordinates, edges are weighted by Euclidean length, main-root vertices *R ⊂ V* form a path rooted at the main-root base *r*_1_, and *L* = *V \R* are lateral-root vertices (Chandrasekhar and Julkowska, 2022). Wiring cost and conduction delay are

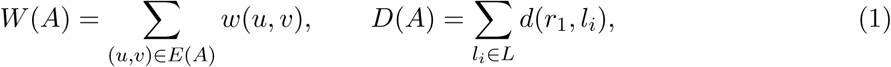

where *d*(*u, v*) is graph distance. Optimal architectures minimize

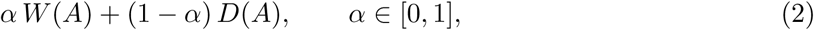

so that *α* = 1 minimizes wiring cost alone and *α* = 0 minimizes conduction delay alone. Holding the main root and the lateral-root tips fixed, each tip is reconnected to the main root along the path that minimizes Eqn 2.

In the straight-line model these paths are line segments. In the curvature-constrained model, a lateral root with tip (*l_x_, l_y_*) connecting to main-root point (*r_x_, r_y_*) follows the quadratic *y*(*x*) = *Gx*^2^+*bx*+*c* passing through both points, so that the single parameter *G* sets the sign and magnitude of curvature (*G >* 0, downward; *G* = 0, straight; *G <* 0, upward; Fig. 1b). A quadratic is the simplest function that represents monotonic curvature with one additional parameter (Altman et al., 2025). For each tip, the optimal attachment point is found by minimizing the arc length of the connecting parabola plus the weighted main-root distance over every admissible main-root segment (Brent’s method; Brent 2013), restricting attachment to main-root segments that existed when the lateral root was initiated (Fig. S2). Derivations are given in Methods S1.

**Figure 1:**
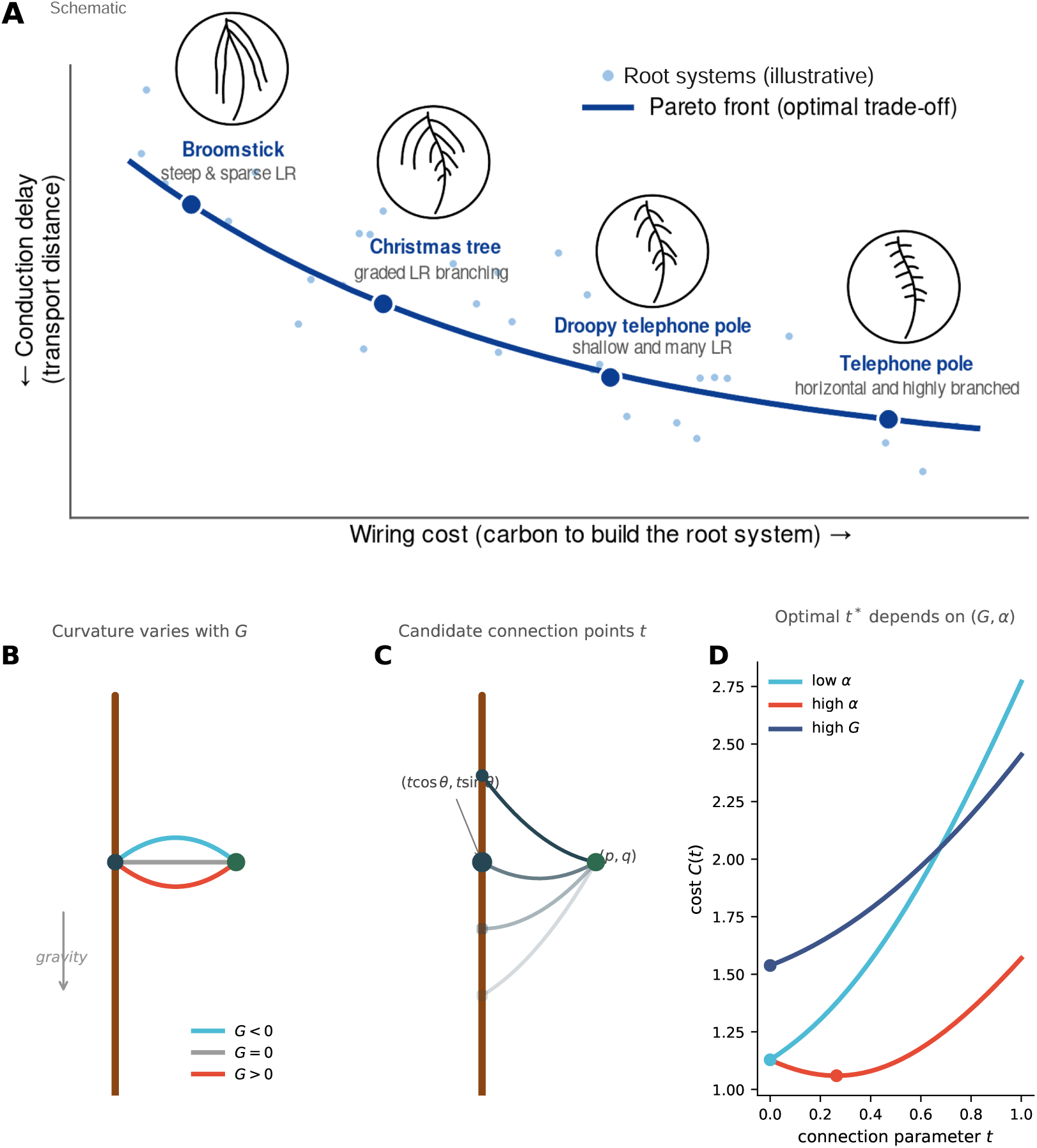
A curvature-constrained Pareto framework for *S. pimpinellifolium* root system architecture. (a) Schematic of the cost–delay trade-off; points are illustrative, not data. Architectures that cannot be improved in one objective without worsening the other form the Pareto front (line). The four architectural types previously described in *S. pimpinellifolium* occupy distinct regions along it. (b) The curvature parameter *G* sets the sign and magnitude of lateral-root curvature (*G <* 0, upward; *G* = 0, straight; *G >* 0, downward). Each lateral root connects to the main root along a quadratic through its tip and its attachment point. (*G, α*) jointly determine the optimal attachment point; the pair that best matches the observed root system is taken as the estimate.

### 2.4 Parameter estimation

We estimated the parameters *G* and *α* for each root system by minimizing the squared orthogonal distance between observed lateral-root points and the corresponding optimal curves (Methods S2, Fig. S1). We first evaluated a coarse grid of parameter combinations, (*G, α*) ∈ [−5, 5] × [0, 1], with step sizes of 1 and 0.1, respectively. We then refined the search around high-performing parameter combinations using finer local grids, including both two-dimensional searches and one-dimensional sweeps along each parameter axis; previously evaluated combinations were excluded from subsequent searches. The parameter pair (*G*, α\**) that minimized the squared orthogonal distance was taken as the estimate. When several grid points tied, we used the mean *G* and the minimum *α* among the tied combinations. Ties affected *G* by more than 0.01 in 0.7% of root systems. The straight-line model was fitted identically, with *G* fixed at 0.

### 2.5 Model comparison

To test whether curvature improves fit beyond the cost of an extra parameter, we treated the squared orthogonal distance as a residual sum of squares and computed, for each root system, ΔBIC = BIC*_G_*_=0_ − BIC_free_ *_G_* with BIC = *n* ln(RSS/*n*) + *k* ln *n*, *k* = 1 (straight) or 2 (curved), and *n* the number of traced lateral-root points. ΔBIC > 2 was taken as support for the curved model.

### 2.6 Direct measurements of root geometry

To validate model parameters against raw geometry, we extracted from the SmartRoot exports, for each root, its insertion angle on the parent root, its arc length, its chord length (base-to-tip vector length) and the direction of that vector. Root angle from vertical was computed from the vector direction (0°, growing straight down; 90°, horizontal), and tortuosity as arc length divided by chord length. Vector direction was exported for PimpiBig5 and PimpiBig6 only (535 root systems); insertion angles were available for all root systems. Per root system, we averaged first-order lateral roots and recorded the main root separately.

### 2.7 Statistical analysis

Analyses were performed in Python with statsmodels (Seabold and Perktold, 2010) and scipy (Virtanen et al., 2020). Salt effects on trait means were estimated with linear mixed models with condition and experiment as fixed effects and accession as a random intercept. Because salt reduces lateral root number and length, and fewer traced points could make parameters less precise, models were fitted both without and with log-transformed number of traced lateral-root points and log-transformed mean lateral-root length as covariates. Salt effects on trait dispersion were tested on the log absolute deviation from the condition median, with the same covariates and accession-clustered standard errors, and confirmed by one-to-one matching of salt and control root systems within experiment on both covariates (Levene’s test, median-centered). Repeatability was estimated as the accession variance share from a random-intercept model within each condition. Differences among architectural types were tested with Kruskal–Wallis tests and with permutation-based PERMANOVA (999 permutations) on standardized parameters (Table S1).

### 2.8 Genome-wide association study

Accession means of the model parameters (*G\**, *α*\* and best-fit distance, for the curved and straight-line models, in each condition; ten traits) were rank-based inverse-normal (INT) transformed. Phenotype codes were linked to genotypes through the accession decode table; 189 accessions had both phenotypes and genotypes. The GWAS phenotypes were computed from root systems imaged on day 5 after transfer, so accessions grown only in PimpiBig5, whose last imaging day was day 4, were not included; the tests of the lead SNP against lateral- and main-root traits, which include PimpiBig5, used up to 231 accessions. Of 17,962,579 SNPs, 5,023,288 passed a minorallele-frequency threshold of 0.05 and a missingness threshold of 0.1, computed on the phenotyped accessions in PLINK (Purcell et al., 2007). Associations were tested in EMMAX (Kang et al., 2010) with an IBS kinship matrix and five principal components as covariates. The genome-wide threshold was Bonferroni-corrected (*P* = 0.05/5,023,288 = 9.95 × 10^−9^); false discovery rates (Benjamini–Hochberg) are reported alongside. Calibration was assessed with QQ plots and the genomic inflation factor *λ*. Linkage disequilibrium (LD) around the lead SNP was computed as *r*^2^ in PLINK, and genes were annotated against the LA2093 reference genome (Wang et al., 2020). Orthology of the candidate gene was assessed by reciprocal best hit with BLASTp (*E <* 10^−10^) between the LA2093 proteome and the *Arabidopsis thaliana* reference proteome (UniProt UP000006548). To ask whether the locus acts through lateral-root development, we tested the lead SNP against accession means of lateral-root number, mean lateral-root length, insertion angle and angle from vertical (each condition; INT-transformed; same kinship and covariates), and refitted *α*\* under control conditions with either lateral-root number or mean lateral-root length added as a covariate, comparing against an unadjusted fit on the same 187 accessions.

### 2.9 Cell-type expression analysis

Because *ISA3* is too sparsely detected in tomato root-tip single-cell data to test for cell-type enrichment, we used a published *Arabidopsis* root single-cell dataset carrying author-assigned cell-type labels (GSE158761; Serrano-Ron et al. 2021). Counts, features and barcodes were taken from the series supplementary files, in which each barcode is annotated with a cell type and a time point. For every gene of interest we counted cells with at least one UMI within each annotated cell type, and compared the proportion of cells detected in the columella/root-cap cluster with the proportion across all 6,658 cells by a one-sided Fisher exact test. Genes with established cell-type restriction were included as positive controls (*SMB*, *BRN2*) and as negative controls (*LBD16*, *LBD29*, *PIN3*, *PIN7*, *SHR*, *WOX5*); the negative controls are strongly depleted in the columella/root-cap cluster, which excludes differences in sequencing depth between clusters as an explanation for enrichment. Analysis scripts are deposited with the rest of the code.

### 2.10 Data and code availability

**Data availability.** All tracing data, metadata, and calculated Pareto fronts required to reproduce the analyses and figures in this study has been archived on Zenodo (DOI: 10.5281/zenodo.21383962). Additional data are available from the corresponding author upon reasonable request.

**Software availability.** The full optimization pipeline and analysis code is publicly available on GitHub and archived on Zenodo (DOI: 10.5281/zenodo.21383534). The software repository contains all scripts required to reproduce the computational model, optimization procedure, and figure generation. Dependencies consist entirely of open-source Python libraries. Original images and SmartRoot tracings of the diversity panel are available from Dryad (https://doi.org/10.5061/dryad.nk98sf826; Rahmati Ishka et al. 2026). The previously published single-cell datasets reanalyzed here are available from NCBI GEO under accessions GSE158761 (*Arabidopsis* root; Serrano-Ron et al. 2021) and GSE212405 (tomato root tip; Canto-Pastor et al. 2024). Root-geometry measurements, hormone-experiment images and tracings, GWAS phenotypes, summary statistics and chromosome 6 locus files, and the GWAS code are available from Zenodo (https://doi.org/10.5281/zenodo.23778702).

## 3 Results

### 3.1 A curvature-constrained Pareto front for lateral roots

To determine whether lateral-root curvature changes where root systems sit in cost–delay space, we generated, for each traced root system, a family of optimal architectures across combinations of curvature *G* and trade-off weight *α* (Fig. 2). Varying *G* at fixed *α* changes the shape of each optimal lateral root (Fig. 2a). Varying *α* at fixed *G* moves the attachment point along the main root (Fig. 2b). Together they produce a layered set of Pareto fronts, one per value of *G*, with curved architectures shifted toward higher cost and longer delay (Fig. 2c). Best-fit optimal architectures reproduced the range of shapes seen in the panel, from upward-curving to strongly downward-curving lateral roots (Fig. 3).

**Figure 2:**
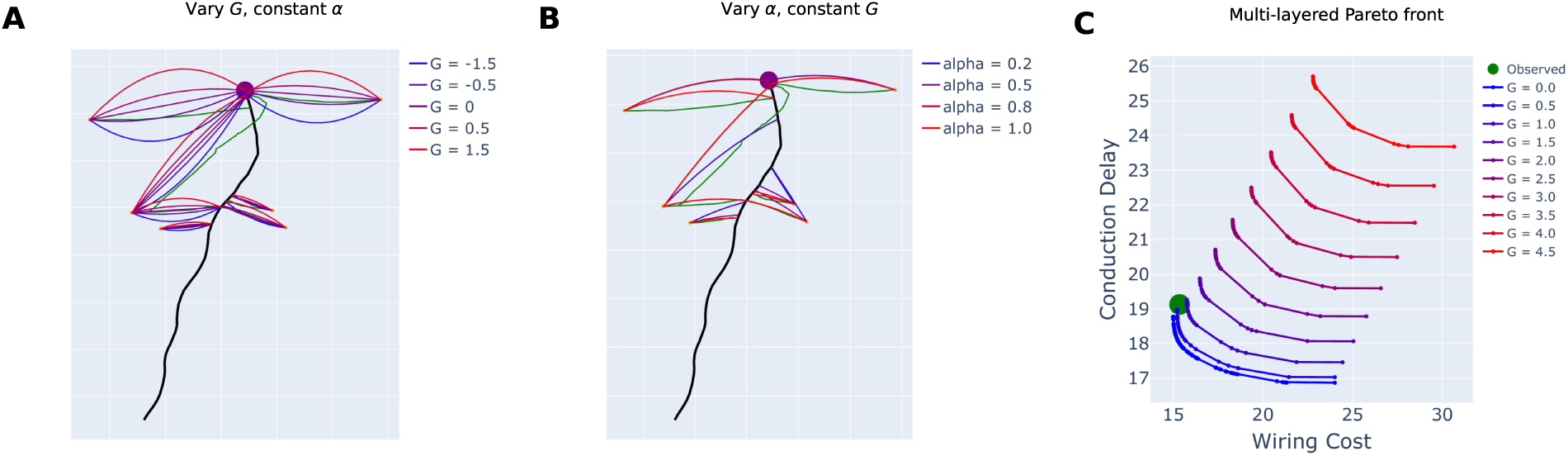
Curvature reshapes the Pareto front of a single root system. (a) Optimal lateral roots at fixed *α* for a range of *G*. (b) Optimal attachment points at fixed *G* for a range of *α*. (c) Pareto fronts for *G* from 0 to 4.5 (colors) and the observed root system (green). Each front is computed with curved lateral roots of the indicated *G*.

**Figure 3:**
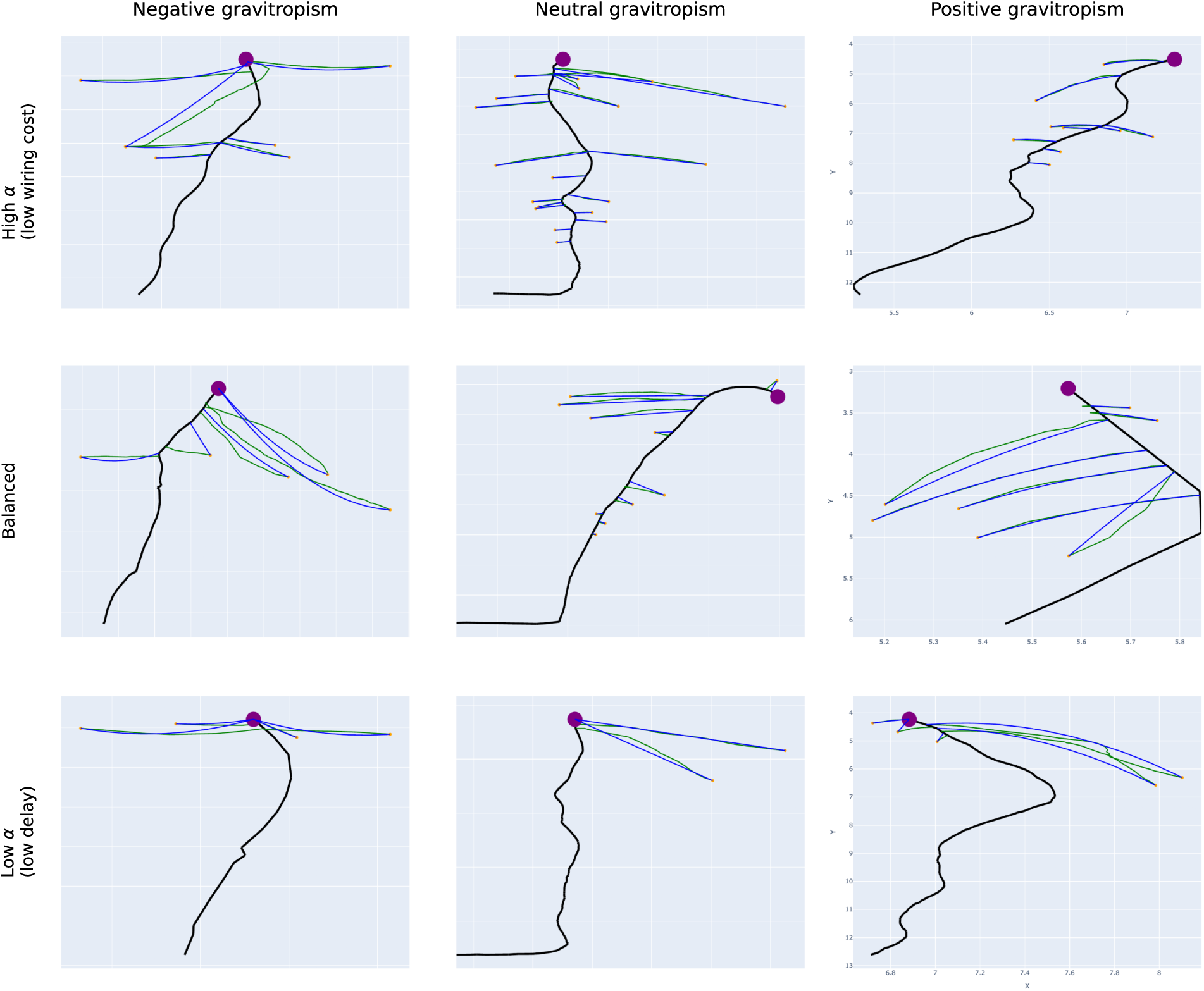
Observed root systems and their best-fit optimal architectures. Observed lateral roots (blue) and best-fit optimal lateral roots (green) for root systems spanning low to high *α* (rows) and negative to positive *G* (columns). Upward curvature corresponds to negative *G*.

### 3.2 Curvature improves the fit beyond the cost of an extra parameter

Because a model with an additional free parameter always fits at least as well as the model nested within it, we compared the curved and straight-line models with a criterion that penalizes the extra parameter. The curved model was supported (ΔBIC > 2) in 80% of control and 75% of salt-grown root systems (median ΔAIC = 79 and 39, respectively), and in 64–91% of root systems within each architectural type and condition (Fig. S4). The curved model never fitted worse than the straight-line model. Allowing curvature also changed the estimated trade-off: *α*\* from the two models was correlated (Spearman *ρ* = 0.85) but differed systematically (Wilcoxon *P <* 10^−80^), in line with the shifts in *α* previously reported by architectural type (Fig. S3). Both parameters differed among architectural types under control conditions (*α*\*, Kruskal–Wallis *H* = 72.9, *P* = 1 × 10^−15^; *G\**, *H* = 87.4, *P* = 8 × 10^−19^), and the two together separated the types under both conditions (PERMANOVA *P* = 0.001; Table S1). Telephone Pole root systems had the lowest curvature and sat at the wiring-cost boundary, whereas Broomstick root systems curved most and had the lowest *α*\*.

To test whether *G* captures real curvature, we compared it with direct measurements from the same tracings. *G\** correlated positively with lateral-root tortuosity (Spearman *ρ* = 0.32 control, 0.22 salt; both *P <* 10^−3^) and negatively with lateral-root angle from vertical (*ρ* = −0.27 and −0.18), but not with insertion angle (*ρ ≈* 0). *G* therefore tracks lateral-root bending, not emergence angle. Its precision, however, depended strongly on how much lateral-root material a root system carried. The absolute deviation of *G\** from the median decreased with the number of traced lateral-root points (*ρ* = −0.31, *P <* 10^−36^; Fig. S5a), and the repeatability of *G\** among replicates of the same accession was low (0.05 control, 0.11 salt; Table 1). We therefore interpret *G* as a curvature correction that improves estimation of *α*, and not as a stand-alone readout of gravitropic sensitivity.

**Table 1:**
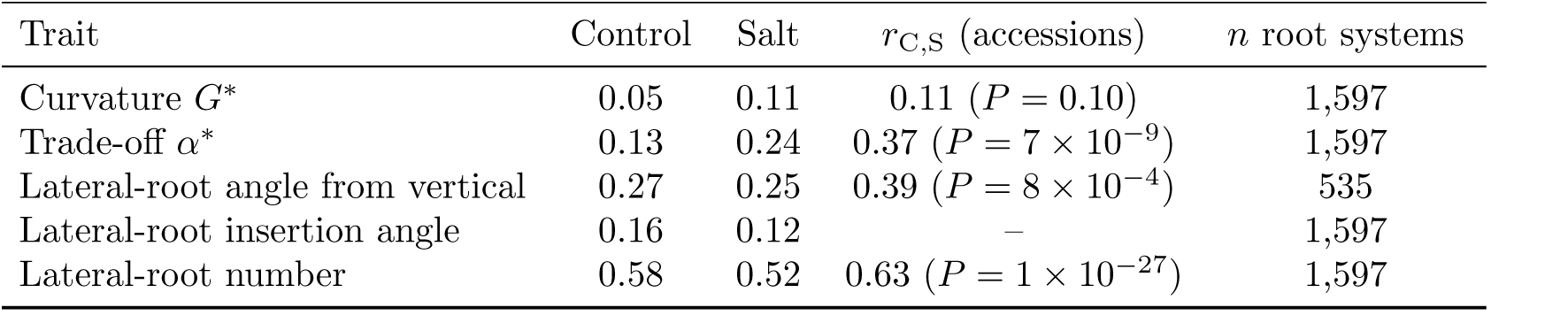
Repeatability of model parameters and direct geometric traits. Accession variance share from random-intercept models within each condition, with experiment as a fixed effect. Genotype-level correlation between conditions is the Pearson correlation of accession means.

### 3.3 Salt shifts root systems toward conduction-delay minimization

Having established that the curved model describes the panel better, we asked how salt stress moves root systems within the trade-off space. Under control conditions most root systems sat at or near the wiring-cost end of the front (mean *α*\* = 0.90; 55% at *α*\* = 1). Salt lowered *α*\* to 0.72 (16% at 1; Fig. 4a). The shift occurred in every architectural type (Mann–Whitney *P ≤* 5 × 10^−9^), and was largest in Broomstick (median 0.92 to 0.39) and Christmas Tree (0.99 to 0.75) root systems. Because salt also reduced lateral-root number (median 17 to 7 per root system) and mean lateral-root length (by 22%), and because *α*\* correlates with both (*ρ* = 0.40 and −0.34 under control conditions), we tested whether the shift simply reflected smaller root systems. It did not: adjusting for the number of traced points and mean lateral-root length, salt still lowered *α*\* by 0.13 (*P* = 6 × 10^−17^; mixed model with accession as random effect). The same shift appeared with the straight-line model (−0.17, *P* = 7 × 10^−23^), so it does not depend on the curvature term. Because *α*\* is a model estimate, we asked whether salt also changed root geometry measured directly from the tracings, independently of the model. Under salt, lateral roots grew closer to vertical: their mean angle from vertical decreased from 66.6° to 56.2° (adjusted difference −10.3°, *P* = 3 × 10^−14^; Fig. 4b), with consistent effects in both experiments where direction was recorded (PimpiBig5, −8.6°, *P* = 0.002; PimpiBig6, −14.0°, *P* = 2 × 10^−7^). Insertion angles also narrowed slightly (72.0° to 68.7°; adjusted −2.9°, *P* = 4 × 10^−5^), and across root systems *α*\* correlated with lateral-root angle from vertical (*ρ* = 0.37 control, 0.43 salt). Because *α* weights cost against transport rather than describing direction, this is a correlation between traits, not a property of the parameter. In contrast, main-root angle did not change (7.4° vs. 6.5° from vertical, *P* = 0.24; Fig. 4c). Salt therefore shifts root systems toward transport efficiency and, in parallel, reorients lateral roots but not the main root.

**Figure 4:**
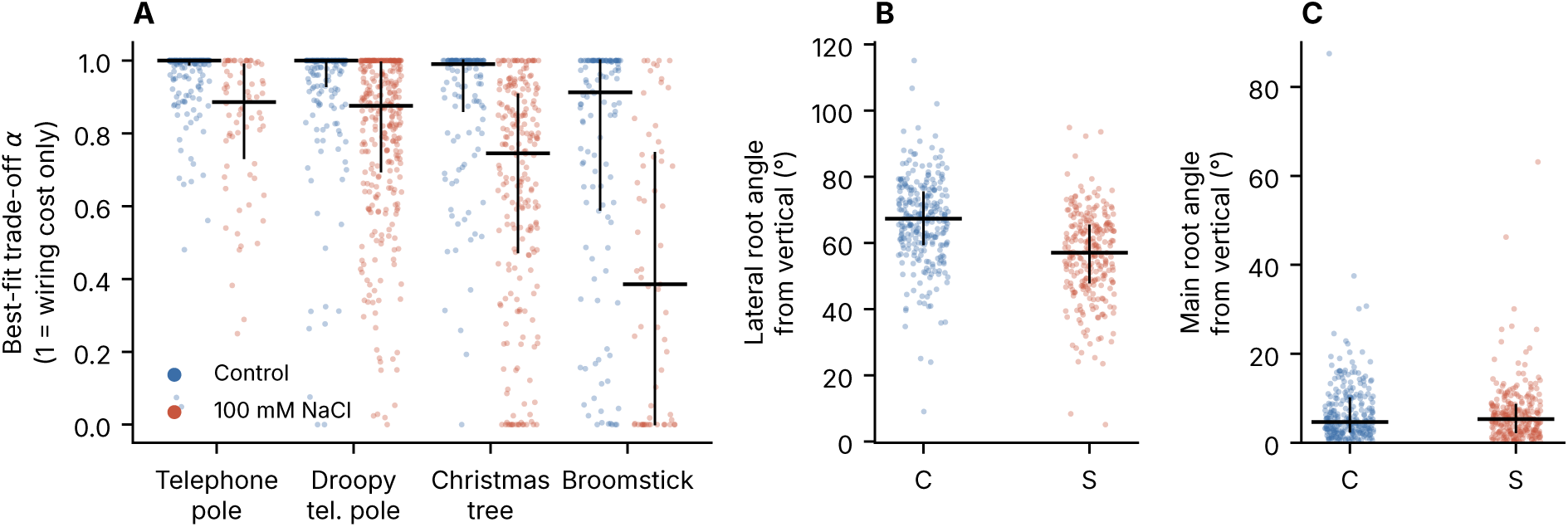
Salt shifts lateral roots, but not the main root, toward transport efficiency. (a) Best-fit trade-off weight *α*\* (curved model) by architectural type under control (blue) and 100 mM NaCl (red). *α* = 1 corresponds to minimizing wiring cost alone. Points are root systems; bars show median and interquartile range. Salt lowered *α*\* in every type (Mann–Whitney *P ≤* 5 × 10^−9^) and after adjustment for lateral-root number and length (mixed model, *P* = 6 × 10^−17^). (b) Mean angle of first-order lateral roots from vertical (0°, straight down), measured directly from tracings (PimpiBig5–6, *n* = 535). Adjusted difference −10.3°, *P* = 3 × 10^−14^. (c) Main-root angle from vertical for the same root systems (*P* = 0.24).

In the raw data, *G\** also appeared more variable under salt (Levene *P* = 7 × 10^−11^). Because *G* is estimated less precisely when lateral-root material is scarce, we tested whether this reflected the smaller root systems. It did: after adjusting for the number of traced points and mean lateral-root length, salt had no effect on the dispersion of *G\** (*P* = 0.65), and in 222 salt–control pairs matched on both covariates the variances did not differ (Levene *P* = 0.45; Fig. S5b). The salt shift in mean *G\** also disappeared after adjustment (*P* = 0.77). Hormone dose–response experiments analyzed with the same covariates are reported in Notes S1.

### 3.4 The trade-off weight is heritable and maps to a locus on chromosome 6

To ask whether the position of a root system on the Pareto front has a genetic basis, we performed a GWAS on accession means of all ten model-derived traits across 189 accessions and 5,023,288 SNPs. SNP-based heritability was modest and consistently higher under salt than under control conditions for seven of eight comparable trait pairs (e.g. *G\**, 0.17 control vs. 0.29 salt; straight-line *α*\*, 0.09 vs. 0.34; Table 2). This agrees with the higher between-replicate repeatability under salt (Table 1). Test statistics were well calibrated for the *G\** and *α*\* traits (*λ* = 0.82–1.01). The three best-fit-distance traits showed inflation (*λ* up to 1.30) without a distinct tail, and we do not interpret them (Fig. S7).

**Table 2:** SNP heritability and calibration of the GWAS for model-derived traits. *h*^2^, EMMAX pseudo-heritability; *λ*, genomic inflation factor. Curved, free-*G* model; straight, *G* = 0 model. Distance traits were inflated and are not interpreted. *^†^*Variance component collapsed to zero; *P*-values for this trait are not kinship-corrected.

| Trait | Model | *h*<sup>2</sup> control | *h*<sup>2</sup> salt | *λ* control | *λ* salt |
|---|---|---|---|---|---|
| *α*<sup>\*</sup> | curved | 0.05 | 0.00<sup>†</sup> | 0.82 | 0.91 |
| *α*<sup>\*</sup> | straight | 0.09 | 0.34 | 0.88 | 0.93 |
| *G*<sup>\*</sup> | curved | 0.17 | 0.29 | 0.95 | 1.01 |
| best-fit distance | curved | 0.24 | 0.33 | 1.30 | 1.15 |
| best-fit distance | straight | 0.26 | 0.31 | 1.14 | 1.03 |

One association exceeded the genome-wide threshold. For *α*\* under control conditions, SNP 6:3,343,493 reached *P* = 1.3 × 10^−9^ (FDR = 0.007; Fig. 5a), supported by neighboring SNPs at 19 and 31 kb (*P ≈* 1 × 10^−7^). The same SNP was the top hit for straight-line *α*\* under control conditions (*P* = 8 × 10^−7^). Because the two traits come from the same plants, this shows the signal does not depend on the curvature term; it is not an independent replication. The minor allele (T; frequency 0.22) lowered *α*\*: median 0.98 in accessions with no copies (*n* = 135), 0.95 in heterozygotes (*n* = 20) and 0.87 in homozygotes (*n* = 33; Fig. 5c). It also reduced the fraction of accessions at the wiring-cost boundary *α*\* = 1 from 26% to 3%. The association remained significant among the 151 accessions below that boundary (*P* = 4.8 × 10^−8^), so it reflects a quantitative shift rather than a boundary artifact. Leave-one-out analysis never raised *P* above 2.2 × 10^−10^, and the lead SNP explained ≈11% of phenotypic variance (an upper estimate, given winner’s curse). As a single pre-specified test, we then examined the lead SNP in the other model-derived traits (Table S2). The T allele also lowered *α*\* under salt (straight-line model, *P* = 1.6 × 10^−4^; curved model, *P* = 1.2 × 10^−3^, not kinship-corrected because the variance component was zero) and increased the distance of salt-grown root systems from their best-fit optimal architecture (straight-line model, the only calibrated distance trait; *P* = 1.5 × 10^−5^). It was not associated with curvature *G\** under either condition (*P* = 0.35 control, 0.39 salt). The locus therefore acts on the wiring–delay trade-off, and not on lateral-root curvature.

**Figure 5:**
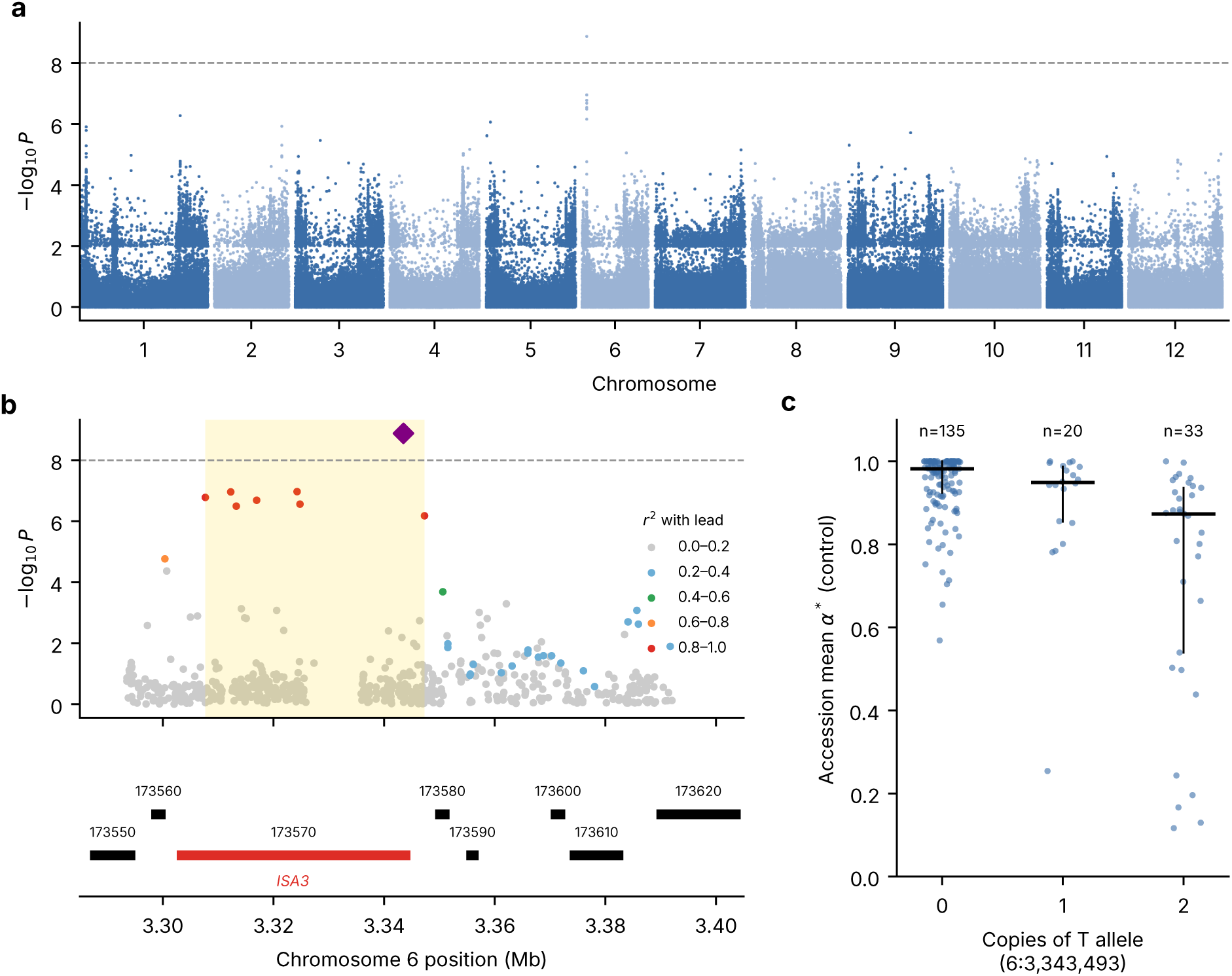
A locus on chromosome 6 associated with the wiring–delay trade-off. (a) Manhattan plot for *α*\* (curved model) under control conditions; 189 accessions, 5,023,288 SNPs, EMMAX with kinship and five principal components. Dashed line, Bonferroni threshold (*P* = 9.95 × 10^−9^). (b) Regional plot of chromosome 6, 3.29–3.40 Mb (shaded: *r*^2^ ≥ 0.8 block), with SNPs colored by *r*^2^ to the lead SNP 6:3,343,493 and gene models below. (c) Accession means of *α*\* (control) by number of copies of the minor (T) allele at the lead SNP; bars, median and interquartile range.

The lead SNP lies within a 39.6-kb LD block (*r*^2^ ≥ 0.8; 3,307,678–3,347,299) containing eight associated SNPs. Seven of them fall in introns of *Spim06g173570* and one lies 1.9 kb from *Spim06g173580* (Fig. 5b). *Spim06g173570* encodes a complete 781-amino-acid protein with all four catalytic regions of glycoside hydrolase family 13 and a predicted plastid transit peptide, and is the reciprocal best BLAST hit of *Arabidopsis* ISOAMYLASE 3 (ISA3, AT4G09020; 71% identity over 738 residues), a starch-debranching enzyme (Wattebled et al., 2005). Its next-best *Arabidopsis* match, ISA1, scored far lower (bit score 592 vs. 1,119), and the two other *S. pimpinellifolium* isoamylase genes are more similar to ISA1 and ISA2, so the assignment to ISA3 is unambiguous. All five common coding variants in *Spim06g173570* (MAF 0.41–0.42) were unassociated with *α*\* (*P* = 0.25–0.85), so the association points to regulatory rather than coding variation. The adjacent gene *Spim06g173580* encodes a cation/H^+^ antiporter (CHX15-like); the lead SNP lies 5.7 kb downstream of its 3*^′^* end.

Because *α*\* correlates with lateral-root number and length, we asked whether the locus acts on lateral-root development and *α*\* merely inherits the effect. The T allele was associated with longer lateral roots under both control (*P* = 3.8 × 10^−6^ at the lead SNP; 3.5 × 10^−7^ within the block) and salt conditions (*P* = 9.5 × 10^−5^), and weakly with fewer lateral roots under control conditions (*P* = 2.3 × 10^−3^), in the directions expected from the correlations of these traits with *α*\* (Table S2). It was not associated with lateral-root insertion angle (*P* = 0.30 control, 0.26 salt; 231 accessions) or angle from vertical (*P ≥* 0.26; 71 accessions). The effect was specific to lateral roots. Main-root length showed at most a weak association in the opposite direction (T allele slightly shorter main roots; *P* = 0.046 control, 0.053 salt; effect less than half that on lateral-root length), and main-root angle was unaffected (*P ≥* 0.34; 71 accessions). Adjusting *α*\* for lateral-root number left the association nearly unchanged (*P* = 1.6 × 10^−8^; effect reduced by 14%). Adjusting for mean lateral-root length weakened it to *P* = 5.5 × 10^−6^ and reduced the effect by 29%, but did not remove it. The locus therefore affects both lateral-root length and the wiring–delay trade-off, and most of its effect on *α*\* is not explained by lateral-root length.

Because the associated variants are non-coding and the cell type in which *ISA3* acts is unknown, we asked where the gene is expressed within the root tip. *ISA3* is too sparsely detected in tomato root-tip single-cell data to support a cell-type test (approximately one transcript per positive cell; Canto-Pastor et al. 2024), so we used a published *Arabidopsis* root single-cell dataset carrying author-assigned cell-type labels (Serrano-Ron et al., 2021). Of the 121 cells in which *ISA3* was detected, 62 fell in the annotated columella/root-cap cluster, which comprises 1,015 of 6,658 cells: a 3.4-fold enrichment (6.1% of columella/root-cap cells versus 1.8% of all cells; one-sided Fisher exact test, *P* = 9.5 × 10^−21^; Fig. S8, Table S4). The enrichment was more specific than that of any other starch-metabolic gene examined: *DPE2* was detected in a larger fraction of columella/root-cap cells but in a larger fraction of cortex cells still, and *AMY3*, *DPE1* and *SEX4* were less strongly enriched. Positive controls behaved as expected (*SMB*, 6.1-fold; *BRN2*, 6.2-fold), whereas markers of other cell types were depleted in the same cluster (*LBD16*, 0.03-fold; *PIN3*, 0.15-fold; *WOX5*, *GATA23* and *LBD29* absent), which excludes greater sequencing depth in that cluster as an explanation.

## 4 Discussion

We set out to make the Pareto description of root architecture geometrically realistic by allowing lateral roots to curve, and to use it to ask how salt moves root systems within the cost–delay trade-off. The curved model fits most root systems better than the straight-line model even after penalizing the extra parameter. With it, salt stress consistently moves *S. pimpinellifolium* root systems away from the wiring-cost end of the front toward conduction-delay minimization. Direct measurements from the same tracings show that salt-grown lateral roots also grow about 10° closer to vertical, whereas the main root is unaffected. The trade-off weight is heritable, more so under salt, and maps to a single locus on chromosome 6.

Allowing curvature matters for estimating the trade-off, but the curvature parameter itself is a weak phenotype. Model comparison with a penalty for the added parameter supported the curved model in three of four root systems, and *α*\* estimates shifted when curvature was allowed, so straight-line estimates of where root systems sit on the front are biased. *G\** correlated with directly measured lateral-root tortuosity and angle, as expected if it captures bending. However, it was estimated imprecisely in root systems with few or short lateral roots and had low repeatability. This imprecision produced an apparent salt-induced increase in the variance of *G*, which disappeared once the amount of lateral-root material was controlled for. We therefore recommend using *G* to correct the geometry of the optimality model, and estimating gravitropic setpoint angles directly from tracings when root direction itself is the question. Per-root curvature parameters, or curvature models with a mechanistic basis in graviperception and proprioception (Bastien et al., 2013), would be natural next steps.

The salt response we describe is organ-specific and directional. Steeper lateral roots under salt are consistent with salt acting on the lateral-root GSA, which in *Arabidopsis* and bean is set by the balance between auxin-driven gravitropic bending and an anti-gravitropic offset (Roychoudhry et al., 2013; Rosquete et al., 2013). Cytokinin contributes to that offset (Waidmann et al., 2019), and salt redistributes auxin in roots (Galvan-Ampudia et al., 2013; Korver et al., 2020). Unlike the main-root agravitropism reported for salt-treated *Arabidopsis* (Sun et al., 2008), main-root angle in *S. pimpinellifolium* was unchanged, which argues that main and lateral roots differ in how salt feeds into directional growth. Lateral roots set their initial growth angle during a short developmental window after emergence, through regulators such as SHOOT GRAVITROPISM 9 that are dispensable for main-root gravitropism (Farkas et al., 2026); salt acting on this lateral-root-specific window would change lateral-root angle without affecting the main root. In the optimality framework, lower *α* means more weight on short transport paths at the cost of more root material. Under salt, where lateral-root growth is restricted, this could favor rapid water delivery from a smaller root system and deeper rooting away from the surface layer where salt accumulates in the field. These interpretations remain to be tested in soil and with genotypes altered in GSA control.

The genetic analysis indicates that position on the Pareto front is partly heritable and that genetic variation is expressed more strongly under salt. Seven of eight trait pairs were more heritable under salt, and repeatability of *α*\* was higher under salt (Table 1). The single significant locus, for *α*\* under control conditions, lies in intronic, non-coding sequence of the *S. pimpinellifolium* ortholog of the starch-debranching enzyme ISA3. The allele that lowers *α*\* also lowers it under salt, places salt-grown root systems further from their best-fit optimal architecture and lengthens lateral roots, but it has no detectable effect on curvature *G\** or on lateral-root insertion angle. Because root gravity sensing depends on starch-filled amyloplasts in columella cells (Kiss et al., 1989), genotypes with larger columella starch granules bend more strongly after reorientation (Sala et al., 2023), and *ISA3* promoter activity is detected in root columella cells (Li et al., 2007), a starch-debranching enzyme could in principle act through graviperception, and the cell-type enrichment reported above places the gene in the right cell type for it (Fig. S8). Even so, the absence of any association with *G\** or insertion angle argues against the graviperception route. A more parsimonious explanation is that the locus acts on carbon allocation: starch turnover governs the carbon available for root growth, and an allele that changes carbon supply to growing lateral roots would alter their length directly. Columella starch itself appears to lie downstream of carbon delivery: in *Arabidopsis* lateral roots, phloem connection to the parent root and the first appearance of columella statoliths both occur within the same 12–24 h window after emergence (Guyomarc’h et al., 2012). The tendency toward shorter main roots in T-allele accessions fits a reallocation of growth from the main root to the laterals rather than a general change in root vigor, and such a reallocation moves the root system along the front toward transport efficiency, consistent with the partial attenuation of the *α*\* association after adjusting for lateral-root length. Because *S. pimpinellifolium* accessions differ across the whole genome, the association does not establish causality. Whether the associated haplotype changes *ISA3* expression, and whether *isa3* loss-of-function alleles alter lateral-root growth and position on the front, remain to be tested.

Our model makes several simplifying assumptions. *G* and *α* are estimated per root system, although individual lateral roots curve differently. Wiring cost and conduction delay are taken as proportional to length, whereas the cost of a root segment depends on its diameter and anatomy, and transport depends on xylem anatomy. Under control conditions, more than half of the root systems were best fit at the wiring-cost boundary (*α*\* = 1), which compresses variation at that end of the front. We constrain *α* to [0, 1] so that the weights assigned to the two objectives in Equation (2) remain nonnegative and sum to one. Negative weights would imply that the objective function rewards worsening one objective, all else equal, which we consider biologically implausible. The clustering of estimates at the boundary value *α*\* = 1 under control conditions could indicate that root architectures under these conditions are best represented by the material-cost extreme of the model. One possibility is that environmental stimuli must exceed a threshold before root systems shift toward architectures that favor faster resource transport. Alternatively, the clustering could reflect limitations of the wiring-cost/conduction-delay trade-off in capturing variation in root system morphology, or differences in the calculation of these objectives across environments that are not explicitly represented in the model. Finally, plants were grown on vertical agar plates, where lateral roots grow along a surface; whether the same trade-off shifts occur in soil remains to be established.

## Acknowledgements

We thank Dr. Chandrasekhar’s former Pitt research assistant Graham Zug, whose observation that “a line *is* a quadratic curve” planted the seed for this work. We thank the C. M. Rick Tomato Genetics Resource Center (University of California, Davis) for seed of *S. pimpinellifolium* accessions, and the Boyce Thompson Institute Computational Biology Center for computing resources.

## Author contributions

AC, MJ and GL conceived the study. AC, AK and MJ developed the model, and KA, AC, AG, AK and KR wrote the software. MJ, MRI and GL performed the plate experiments, and MJ and MRI traced the root systems and curated the data. KA, AC, AK, MJ, MRI and KR analyzed the model fits. RN and MJ performed the GWAS and the reanalysis of root geometry. AC and MJ wrote the manuscript with input from all authors. AC, MJ and KR supervised the work.

## AI usage statement

Generative AI tools (ChatGPT, OpenAI; Claude, Anthropic) were used for code generation, debugging, data quality control, exploratory and statistical analysis, visualization and editing of text. All outputs were reviewed, validated and, where necessary, modified by the authors, who take full responsibility for the content.

