## Supplemental Information for "Salt stress shifts wild tomato root systems toward transport efficiency along a gravitropism-constrained Pareto front"

#### Methods S1. Constructing an optimal architecture

Our previous mathematical model constructs an architecture that minimizes

$$C(t) = \alpha \cdot W(A) + (1 - \alpha) \cdot D(A) \tag{S1}$$

where $W(.)$ is the wiring cost and $D(.)$ is the conduction delay between the lateral roots and the main root. By considering only these objectives, the algorithm was incentivized to connect each lateral root tip to the main root via a straight line. We will extend the algorithm to take as input a gravitropic constant $G$.

With the introduction of a gravitropic constant, $G$, we are able to model a plant's tendency to grow towards the gravity axis. If $G = 0$, then the lateral root connects using straight lines. Therefore, the larger $G$ is, the more quadratic the curve of the lateral root will be. Given a main root anchored at the origin $(0, 0)$ with length $l$ and angle of inclination $\theta$, the lateral root starts on the main root at the point $(tlcos\theta, tlsin\theta)$ $(t \in [0, 1])$ and ends at the point $(p, q)$

To represent the lateral root as a curve, we use the equation

$$y(x) = Gx^2 + b(t)x + c(t) \tag{S2}$$

where G is the quadratic coefficient, b is the linear coefficient, and c is the constant coefficient. It follows that the modeled lateral root interpolates the points $(tlcos\theta, tlsin\theta)$ and $(p, q)$ when

$$b(t) = \frac{G(t\ell\cos\theta)^2 - t\ell\sin\theta + q - Gp^2}{p - t\ell\cos\theta}, \text{ and}$$

$$c(t) = \frac{p(t\ell\sin\theta - G(t\ell\cos\theta)^2) - t\ell\cos\theta(q - Gp^2)}{p - t\ell\cos\theta},$$

with $t\ell\cos\theta \neq p$.

Given $y(x) = Gx^2 + b(t)x + c(t)$, we then need to calculate the length of the curve from $(tlcos\theta, tlsin\theta)$ to $(p, q)$. From calculus, the formula for this is:

$$L(t) = \int_{tlcos\theta}^{p} \sqrt{1 + (y'(x))^2}dx = \int_{tlcos\theta}^{p} \sqrt{1 + (2Gx + b(t))^2}dx$$

.

We seek to find the value of $t$ that minimizes

$$\begin{aligned} C(t) &= \alpha \cdot W(t) + (1-\alpha) \cdot D(t) \\ &= \alpha \cdot L(t) + (1-\alpha) \cdot (L(t) + tl) \\ &= L(t) + (1-\alpha) \cdot tl \\ &= \left( \int_{tlcos\theta}^{p} \sqrt{1 + (2Gx + b(t))^2} dx \right) + (1-\alpha) \cdot tl \end{aligned} \tag{S3}$$

**Approach 1:** Using calculus, one can show that:

$$L(t) = \int_{tlcos\theta}^{p} \sqrt{1 + (2Gx + b(t))^2} dx = \frac{1}{2G} \left[ \frac{1}{2} \sec\theta \tan\theta + \frac{1}{2} \ln \left| \sec\theta + \tan\theta \right| \right] \Bigg|_{\theta_0(t)}^{\theta_p(t)}$$

Where

$$\begin{aligned} \theta_0(t) &= \arctan\left( \frac{q - Gp^2 - t}{p} \right) \\ \theta_p(t) &= \arctan\left( 2Gp + \frac{q - Gp^2 - t}{p} \right). \end{aligned}$$

See (Altman et al., 2025) for partial derivation. We can then re-write equation (S1) as

$$\begin{aligned} C(t) &= \left( \int_{tlcos\theta}^{p} \sqrt{1 + (2Gx + b(t))^2} dx \right) + (1-\alpha) \cdot tl \\ &= \frac{1}{2G} \left[ \frac{1}{2} \sec\theta \tan\theta + \frac{1}{2} \ln \left| \sec\theta + \tan\theta \right| \right] \Bigg|_{\theta_0(t)}^{\theta_p(t)} + (1-\alpha) \cdot tl \end{aligned} \tag{S4}$$

Equation S4 defines a single variable function on a restricted domain. In practice, this function is also often convex. Thus, the global minimum can be efficiently approximated using approaches such as Brent's method (Brent, 2013), which is implemented in many modern numerical analysis software packages (including Scipy Virtanen et al. 2020).

**Approach 2:** We are aiming to minimize equation (S1), so therefore we must minimize the following

$$C(t) = \alpha \cdot L(t) + (1-\alpha) \cdot (L(t) + tl) \tag{S5}$$

Which simplifies down to

$$C(t) = \int_{tlcos\theta}^{p} \sqrt{1 + (2Gx + b(t))^2} dx + (1-\alpha)tl.$$

.

Using Liebniz's Integral Rule, we have

$$\begin{aligned}
L'(t) &= \frac{d}{dt}\int_{t\ell\cos\theta}^{p}\left(\sqrt{1+(2Gx+b(t))^2}\right)dx - \left(\sqrt{1+(2Gx+b(t))^2}\right|_{x=t\ell\cos\theta}\cdot\ell\cos\theta \\
&= \int_{t\ell\cos\theta}^{p}\frac{\partial}{\partial t}\left(\sqrt{1+(2Gx+b(t))^2}\right)dx - \left(\sqrt{1+(2Gx+b(t))^2}\right|_{x=t\ell\cos\theta}\cdot\ell\cos\theta \\
&= \int_{t\ell\cos\theta}^{p}\frac{(2Gx+b(t))b'(t)}{\sqrt{1+(b(t)+2Gx)^2}}dx - \ell\cos\theta\sqrt{1+(2Gt\ell\cos\theta+b(t))^2} \\
&= \frac{b'(t)}{2G}\int_{t\ell\cos\theta}^{p}\frac{(2Gx+b(t))2G}{\sqrt{1+(2Gx+b(t))^2}}dx - \ell\cos\theta\sqrt{1+(2Gt\ell\cos\theta+b(t))^2} \\
&= \frac{b'(t)}{2G}\int_{t\ell\cos\theta}^{p}\frac{\partial}{\partial x}\left(\sqrt{(2Gx+b(x))^2+1}\right)dx - \ell\cos\theta\sqrt{1+(2Gt\ell\cos\theta+b(t))^2} \\
&= \frac{b'(t)}{2G}\left(\sqrt{1+(2Gp+b(t))^2}-\sqrt{1+(2Gt\ell\cos\theta+b(t))^2}\right) - \ell\cos\theta\sqrt{1+(2Gt\ell\cos\theta+b(t))^2} \\
&= \frac{b'(t)}{2G}\sqrt{1+(2Gp+b(t))^2} - \left(\ell\cos\theta+\frac{b'(t)}{2G}\right)\sqrt{1+(2Gt\ell\cos\theta+b(t))^2}
\end{aligned}$$

Thus $C'(t) = L'(t) + (1-\alpha)l = 0$ when

$$\frac{b'(t)}{2G}\sqrt{1+(2Gp+b(t))^2} - \left(\ell\cos\theta+\frac{b'(t)}{2G}\right)\sqrt{1+(2Gt\ell\cos\theta+b(t))^2} - (1-\alpha)\ell = 0. \quad \text{(S6)}$$

When $\theta = \frac{\pi}{2}$ (i.e., the main root is completely vertical), one can derive an analytical solution to equation (S6) (Altman et al., 2025). In the general case, equation (S6) can be efficiently and accurately solved numerically using standard Python packages in the SciPy library (Virtanen et al., 2020). If the roots of the equation lie $\in [0,1]$, then the connection point on the main root is determined by using $(tlcos\theta, tlsin\theta)$. Otherwise, the endpoints of the main root are tested and the lower costing path is chosen.

#### Generalizing to multiple lateral root tips and main root segments

The framework developed above assumes a single lateral root tip and a single main root segment anchored at the origin. We now generalize to an arbitrary root architecture in which the main root is a polygonal chain of $M$ segments and the plant has $N$ lateral root tips.

Each lateral root tip $i$ can be optimized independently of all others, since the cost function decomposes additively across tips. For a given tip $(p_i, q_i)$, we find the optimal connection point by optimizing over all main root segments jointly. For segment $j$, with base located at cumulative distance $d_j$ from the root base, the total cost acquires an additive penalty $d_j$ reflecting the signal propagation distance along the main root prior to reaching that segment. The optimal connection for tip $i$ is then:

$$(t_i^*, j_i^*) = \underset{\substack{j\in\{1,\ldots,M\} \\ t\in[0,1]}}{\arg\min}\ [\alpha\cdot w(t,j,p_i,q_i) + (1-\alpha)\,(w(t,j,p_i,q_i) + d_j + t\ell_j)] \quad \text{(S7)}$$

where $w(t, j, p_i, q_i)$ denotes the arc length of the optimal parabola connecting the branch point on segment $j$ at parameter $t$ to tip $(p_i, q_i)$, $\ell_j$ is the length of segment $j$, and $d_j$ is the cumulative main root length from the base to the start of segment $j$. The optimal branch point on each segment is found using Brent's method (Brent, 2013), and the globally optimal segment $j_i^*$ and position $t_i^*$ are determined by taking the minimum over all segments. The total cost for the arbor is then

$$\sum_{i=1}^{N} \text{cost}(t_i^*, j_i^*, p_i, q_i)$$

#### Methods S2. Comparing root systems by squared orthogonal distance

To determine values $(G^*, \alpha^*)$ that best represent the structure of a given root system, we need a metric that quantifies structural similarity between an observed architecture and the optimal architecture generated by a parameter setting $(G, \alpha)$. Earlier models compared observed and optimal architectures by their wiring costs and conduction delays in Pareto-front space (Chandrasekhar and Navlakha, 2019; Chandrasekhar and Julkowska, 2022). This fails for a curved model, because two curves with different geometry can share the same wiring cost and conduction delay (Fig. S1a). Instead, we quantify how closely the shapes of optimal lateral roots follow the shapes of their observed counterparts.

We use the *squared orthogonal distance*, the squared perpendicular distance from each observed point to the optimal curve (Fig. S1b). Both the $x$- and $y$-coordinates of traced points carry measurement error, because the tracer sets both coordinates jointly. Minimizing vertical residuals alone, as in ordinary least squares, would treat the $x$-coordinates as error-free. Orthogonal distance regression treats both coordinates symmetrically (Fuller, 1987; Boggs et al., 1989).

Let $l$ be a lateral root comprising a set of traced segments. Each segment is sub-discretized into 100 evenly spaced points, giving a point set $\{(x_1, y_1), \ldots, (x_n, y_n)\}$ for that lateral root, so that lateral roots are treated consistently regardless of tracing density. For an optimal lateral root following the quadratic $f_l$, the distance is

$$D(l, f_l) = \sum_i \min_x \left[(x - x_i)^2 + (f_l(x) - y_i)^2\right], \tag{S8}$$

where the minimum is over all points on $f_l$. For the set of lateral roots $L = \{l_1, \ldots, l_k\}$ and optimal curves $F = \{f_{l_1}, \ldots, f_{l_k}\}$, the total distance is

$$\mathcal{D}(L, F) = \sum_i D(l_i, f_{l_i}), \tag{S9}$$

and

$$(G^*, \alpha^*) = \underset{G,\alpha}{\arg\min}\ \mathcal{D}(L, F_{G,\alpha}), \tag{S10}$$

where $F_{G,\alpha}$ is the set of optimal lateral-root curves under $(G, \alpha)$. [CHECK: Arjun: confirm this squared form is what best_sq_orthogonal.csv uses.]

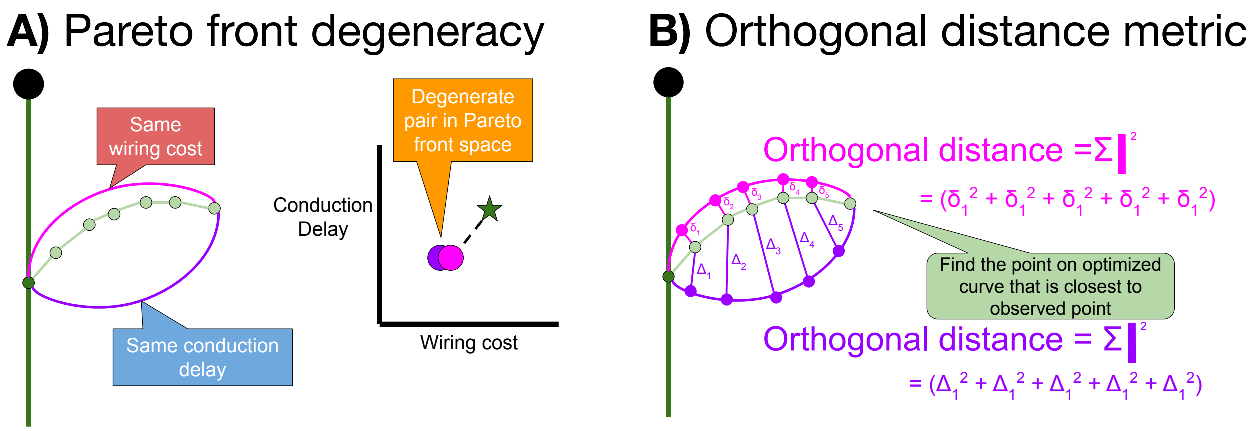


Figure S1: **Squared orthogonal distance.** (a) Distinct optimal lateral roots can have identical Pareto costs although they differ in how closely they follow the observed root. (b) Similarity is therefore measured as the sum of squared orthogonal distances from discretized points along the observed root to the optimal curve.

#### Methods S3. Constraint on attachment points

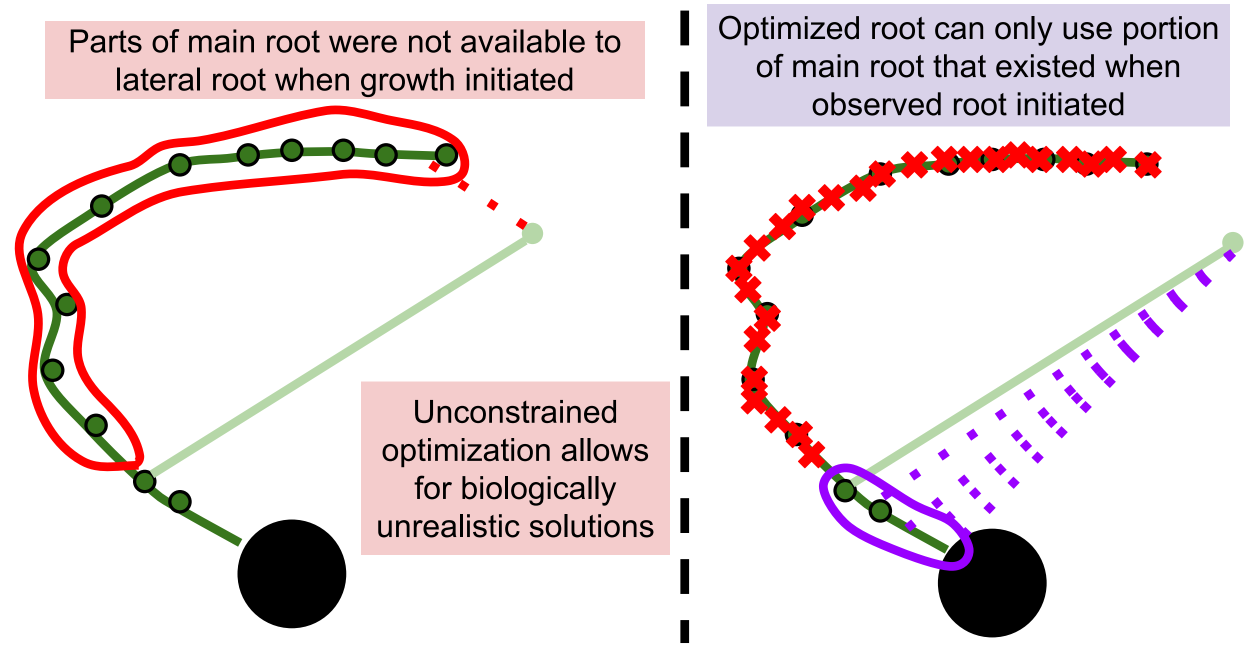


Figure S2: **Constraint on lateral root attachment points.** Observed lateral roots can only attach to portions of the main root that existed at the time of their initiation. We impose the same constraint on optimized lateral roots to prevent biologically unrealistic connections to segments that arise later in development.

### Table S1. Differences among architectural types

Table S1: **Differences in model parameters among architectural types.** Kruskal–Wallis tests on $\alpha^*$ and $G^*$, and PERMANOVA (Euclidean distance on standardized parameters, 999 permutations) on each parameter and on both jointly. Diversity-panel root systems with one of the four type labels, squared-distance fits: $n = 706$ (control) and 679 (salt).

| Condition | Test | Model | Statistic | $P$ |
|---|---|---|---|---|
| Control | Kruskal–Wallis | $\alpha \sim$ type | $H = 72.9$ | $1.0 \times 10^{-15}$ |
| Control | Kruskal–Wallis | $G \sim$ type | $H = 87.4$ | $8.1 \times 10^{-19}$ |
| Control | PERMANOVA | $\alpha \sim$ type | pseudo-$F = 36.0$ | 0.001 |
| Control | PERMANOVA | $G \sim$ type | pseudo-$F = 5.0$ | 0.001 |
| Control | PERMANOVA | $(G, \alpha) \sim$ type | pseudo-$F = 19.6$ | 0.001 |
| Salt | Kruskal–Wallis | $\alpha \sim$ type | $H = 86.1$ | $1.5 \times 10^{-18}$ |
| Salt | Kruskal–Wallis | $G \sim$ type | $H = 18.0$ | $4.4 \times 10^{-4}$ |
| Salt | PERMANOVA | $\alpha \sim$ type | pseudo-$F = 43.7$ | 0.001 |
| Salt | PERMANOVA | $G \sim$ type | pseudo-$F = 2.2$ | 0.11 |
| Salt | PERMANOVA | $(G, \alpha) \sim$ type | pseudo-$F = 21.2$ | 0.001 |

#### Supporting figures

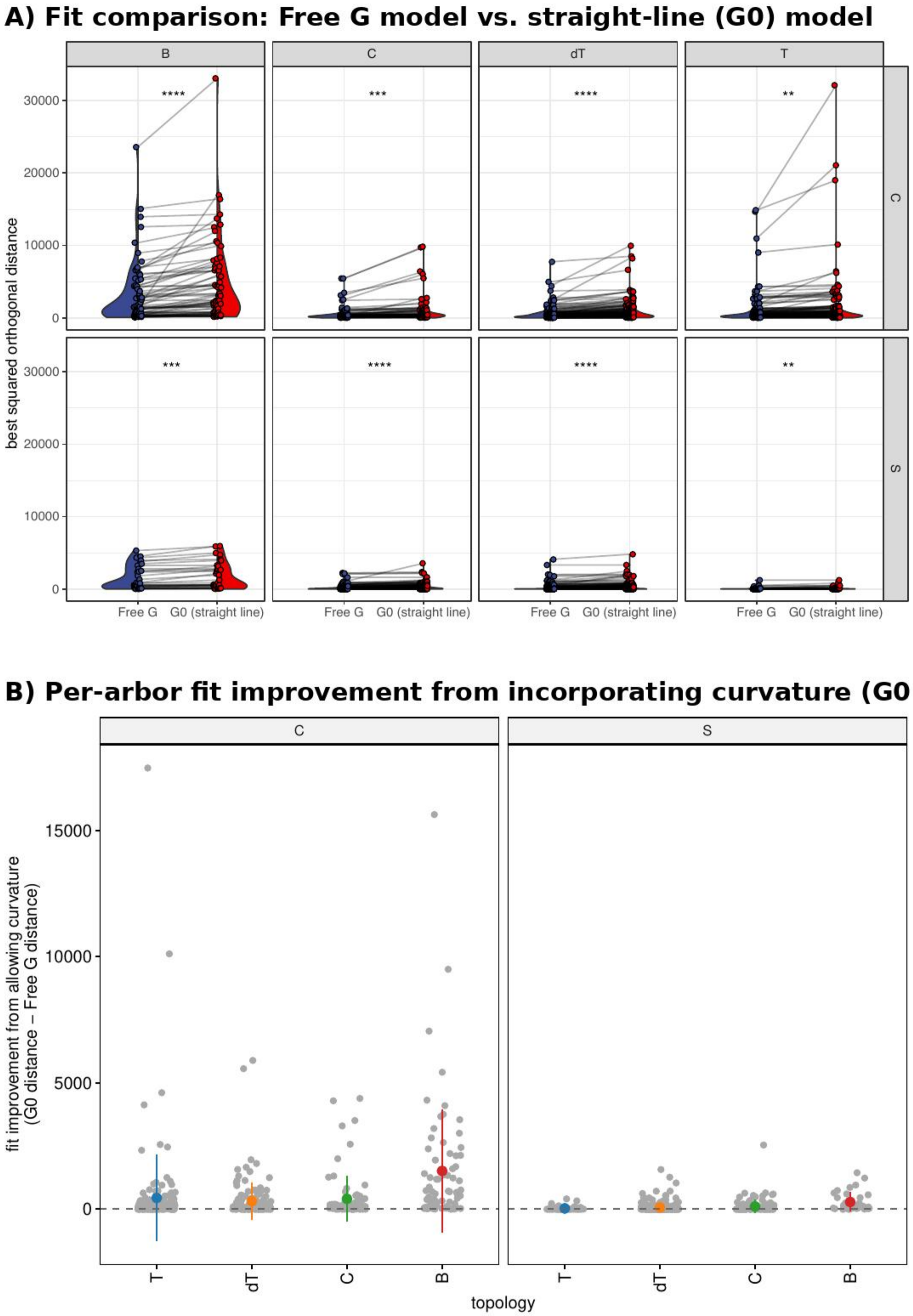


Figure S3: **Fit of the curved (free $G$) and straight-line ($G = 0$) models.** (a) Best-fit squared orthogonal distance per root system under free $G$ (blue) and $G = 0$ (red), by architectural type (Telephone Pole, T; Christmas Tree, C; Droopy Telephone Pole, dT; Broomstick, B) and condition (control, C; salt, S). Lines connect the same root system. Free $G$ never fits worse than $G = 0$; Fig. S4 gives the penalized comparison (Wilcoxon signed-rank; $^{**}P<0.01$, $^{***}P<0.001$, $^{****}P<0.0001$). (b) Per-root-system fit improvement (distance under $G = 0$ minus distance under free $G$). Improvement is larger under control conditions, where root systems carry more lateral-root material (Fig. S5).

Figure not uploaded yet:
`Figs/Model-Comparison/FigS_model_comparison_BIC.pdf`

Figure S4: **Penalized comparison of the curved and straight-line models.** Per root system, $\Delta\text{BIC} = \text{BIC}_{G=0} - \text{BIC}_{\text{free}\,G}$ (signed $\log_{10}(1 + |\Delta|)$ scale), by architectural type, control (blue) and salt (red). Dashed line, $\Delta\text{BIC} = 2$. The curved model was supported in 80% (control) and 75% (salt) of root systems.

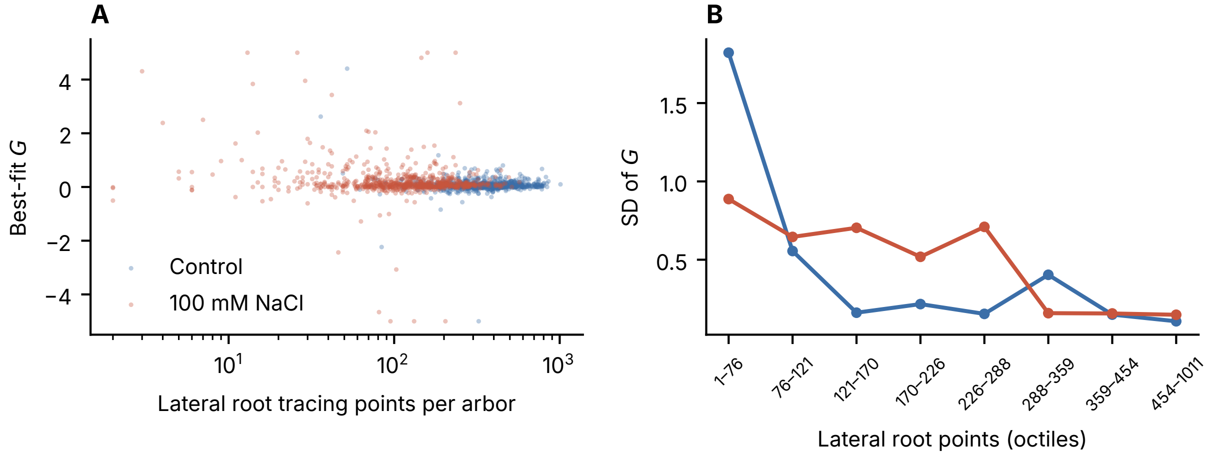


Figure S5: **Precision of $G^*$ depends on the amount of lateral-root material.** (a) $G^*$ against the number of traced lateral-root points per root system. (b) Standard deviation of $G^*$ within octiles of traced points. Salt-grown root systems are concentrated in the low-data octiles; within octiles, salt and control spreads are similar (adjusted dispersion test $P = 0.65$; matched pairs, Levene $P = 0.45$).

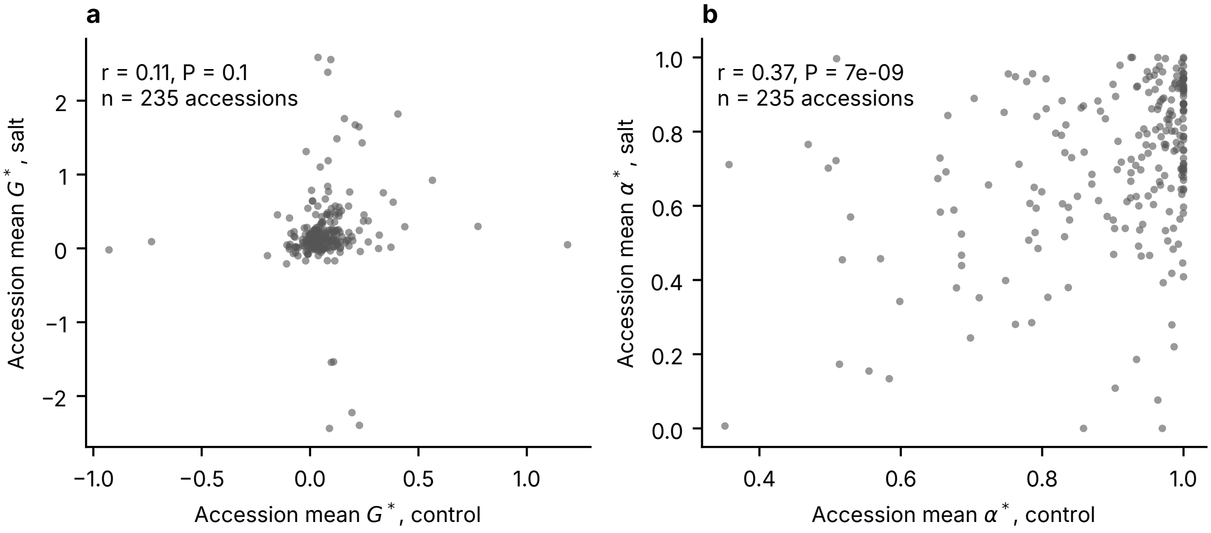


Figure S6: **Accession-level correlations between control and salt conditions.** (a) $G^*$, $r = 0.11$, $P = 0.10$. (b) $\alpha^*$, $r = 0.37$, $P = 7 \times 10^{-9}$. Points are accession means across replicates within each condition; 235 accessions with data in both conditions.

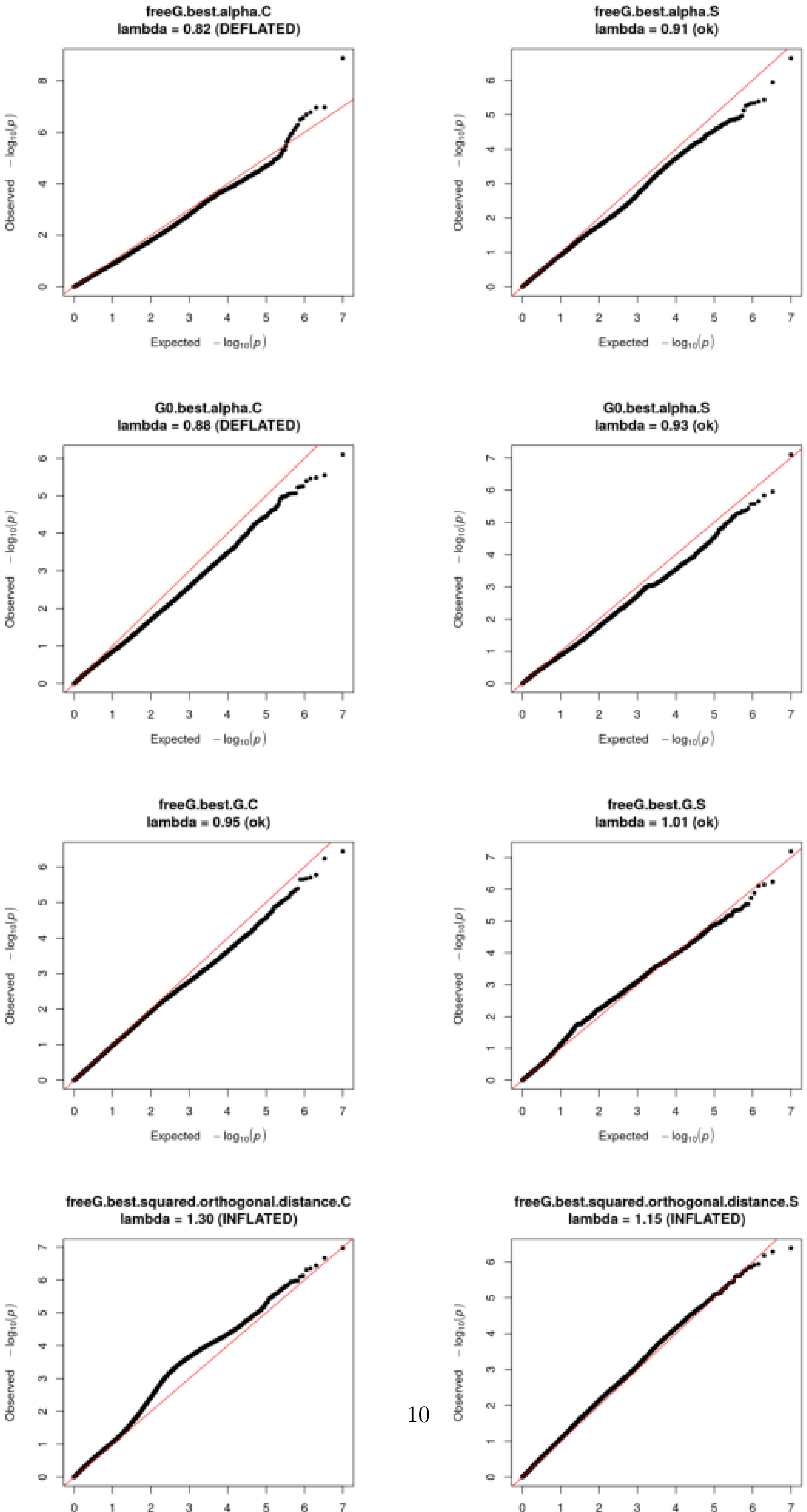
freeG.best.alpha.C
lambda = 0.82 (DEFLATED)
freeG.best.alpha.S
lambda = 0.91 (ok)
G0.best.alpha.C
lambda = 0.88 (DEFLATED)
G0.best.alpha.S
lambda = 0.93 (ok)
freeG.best.G.C
lambda = 0.95 (ok)
freeG.best.G.S
lambda = 1.01 (ok)
freeG.best.squared.orthogonal.distance.C
lambda = 1.30 (INFLATED)
freeG.best.squared.orthogonal.distance.S
lambda = 1.15 (INFLATED)
Expected $-\log_{10}(p)$
Observed $-\log_{10}(p)$

### Table S2. Chromosome 6 lead SNP across model parameters and root traits

Table S2: **Association of the chromosome 6 lead SNP (6:3,343,493) with model parameters and lateral- and main-root traits, and conditional tests for $\alpha^*$.** EMMAX on chromosome 6 with the genome-wide kinship matrix and five principal components; phenotypes are INT-transformed accession means. $\beta$ is for the coded allele (the major allele; the minor T allele has the opposite sign). Best $P$ in block, lowest $P$ within the $r^2 \geq 0.8$ block (3,307,678–3,347,299). Conditional fits use the same 187 accessions as the unadjusted fit. Rows marked ‡ are read from the genome-wide run (same kinship and covariates); they are single tests of one pre-specified SNP. †Variance component collapsed to zero; not kinship-corrected.

| Trait | Accessions | $\beta$ (lead) | $P$ (lead) | Best $P$ in block |
|---|---|---|---|---|
| $\alpha^*$, control (unadjusted, same subset) | 187 | 0.55 | $8.5 \times 10^{-10}$ | $8.5 \times 10^{-10}$ |
| $\alpha^*$, control, adjusted for lateral-root number | 187 | 0.48 | $1.6 \times 10^{-8}$ | $1.6 \times 10^{-8}$ |
| $\alpha^*$, control, adjusted for mean lateral-root length | 187 | 0.39 | $5.5 \times 10^{-6}$ | – |
| $\alpha^*$, salt, straight-line model‡ | 188 | 0.38 | $1.6 \times 10^{-4}$ | $1.6 \times 10^{-4}$ |
| $\alpha^*$, salt, curved model‡† | 188 | 0.33 | $1.2 \times 10^{-3}$ | $1.2 \times 10^{-3}$ |
| Best-fit distance, salt, straight-line model‡ | 188 | −0.40 | $1.5 \times 10^{-5}$ | $1.5 \times 10^{-5}$ |
| Curvature $G^*$, control‡ | 188 | −0.09 | 0.35 | 0.040 |
| Curvature $G^*$, salt‡ | 188 | −0.09 | 0.39 | 0.009 |
| Mean lateral-root length, control | 231 | −0.40 | $3.8 \times 10^{-6}$ | $3.5 \times 10^{-7}$ |
| Mean lateral-root length, salt | 228 | −0.34 | $9.5 \times 10^{-5}$ | $5.2 \times 10^{-5}$ |
| Lateral-root number, control | 231 | 0.26 | $2.3 \times 10^{-3}$ | $3.4 \times 10^{-4}$ |
| Lateral-root number, salt | 228 | 0.04 | 0.67 | $5.4 \times 10^{-3}$ |
| Lateral-root insertion angle, control | 231 | 0.09 | 0.30 | $4.0 \times 10^{-3}$ |
| Lateral-root insertion angle, salt | 228 | 0.11 | 0.26 | $1.3 \times 10^{-4}$ |
| Lateral-root angle from vertical, control | 71 | −0.19 | 0.26 | 0.036 |
| Lateral-root angle from vertical, salt | 70 | 0.09 | 0.59 | 0.043 |
| Main-root length, control | 231 | 0.18 | 0.046 | – |
| Main-root length, salt | 228 | 0.17 | 0.053 | – |
| Main-root angle from vertical, control | 71 | −0.17 | 0.34 | – |
| Main-root angle from vertical, salt | 70 | 0.01 | 0.95 | – |

### Notes S1. Hormone dose–response experiments

To ask whether hormone treatments alter lateral-root curvature, we applied the model to root systems from dose-controlled experiments with two *S. pimpinellifolium* accessions, LA1371 (KAUST M058) and LA2540 (M248) (Ishka et al., 2026): abscisic acid (ABA; 0, 1, 10 µM), the ethylene precursor ACC (0, 1, 5 µM), gibberellic acid ($GA_3$; 0, 1, 5 µM), trans-zeatin (0, 1, 10 µg mL$^{-1}$) and IAA (0 or 0.1 µM; a 0.05 µM treatment in the same experiment was not traced), each with and without 100 mM NaCl, imaged on day 9. The cultivated accession LA1511 (*S. lycopersicum*) from the same experiments was excluded. Because the precision of $G$ depends on the amount of lateral-root material (Fig. S5) and hormones change lateral-root number, all models included log number of traced lateral-root points and log lateral-root number as covariates, with accession as a fixed effect. For each hormone we fitted $G^* \sim$ salt $\times$ dose for the mean and the same model on log absolute deviations from group medians for dispersion (type II ANOVA; Table S3).

After adjustment, two effects survived correction for the 30 tests in the table (Bonferroni $P <$ 0.0017). In the ACC experiment, salt lowered mean $G^*$ (more upward or less downward curvature;

$P = 8 \times 10^{-5}$), with no dose effect. In the trans-zeatin experiment, the dispersion of $G^*$ depended on the combination of salt and dose ($P = 8 \times 10^{-4}$). IAA reduced mean $G^*$ and its dispersion at nominal significance only ($P = 0.015$ and 0.002). Several effects reported in an earlier version of this analysis without lateral-root covariates (salt effect on $G$ variance under ACC; dose effect on $G$ variance under trans-zeatin, independent of salt) were not robust to adjustment. Across the hormone experiments, salt lowered $\alpha^*$ ($-0.16$, $P = 2 \times 10^{-14}$), and the effect was attenuated but remained after adjusting for lateral-root amount ($-0.07$, $P = 0.005$; non-ACC experiments, where lateral-root length was available), in the same direction as in the diversity panel. With two accessions and 13–20 root systems per group, these experiments have limited power, and we treat them as exploratory.

Table S3: **Hormone effects on $G^*$ after adjusting for lateral-root amount.** *S. pimpinellifolium* accessions LA1371 (M058) and LA2540 (M248). $F$ and $P$ from type II ANOVA of $G^* \sim$ salt $\times$ dose + accession + log(traced points) + log(lateral-root number) (mean) and of the log absolute deviation from the group median with the same terms (dispersion). Bonferroni threshold for 30 tests, $P = 0.0017$.

| Hormone | Term | $n$ | Mean $F$ | Mean $P$ | Dispersion $F$ | Dispersion $P$ |
|---|---|---|---|---|---|---|
| ABA | salt | 113 | 1.3 | 0.25 | 3.4 | 0.068 |
| ABA | dose | 113 | 2.2 | 0.11 | 3.1 | 0.048 |
| ABA | salt $\times$ dose | 113 | 0.8 | 0.43 | 1.4 | 0.25 |
| ACC | salt | 118 | 16.8 | $8.1 \times 10^{-5}$ | 0.5 | 0.47 |
| ACC | dose | 118 | 0.6 | 0.57 | 0.1 | 0.87 |
| ACC | salt $\times$ dose | 118 | 0.3 | 0.75 | 0.2 | 0.78 |
| $GA_3$ | salt | 119 | 0.7 | 0.39 | 1.7 | 0.2 |
| $GA_3$ | dose | 119 | 1.7 | 0.18 | 4.5 | 0.013 |
| $GA_3$ | salt $\times$ dose | 119 | 0.9 | 0.41 | 0.9 | 0.41 |
| trans-zeatin | salt | 108 | 3.0 | 0.089 | 0.0 | 0.86 |
| trans-zeatin | dose | 108 | 1.7 | 0.19 | 5.4 | 0.0062 |
| trans-zeatin | salt $\times$ dose | 108 | 0.9 | 0.42 | 7.7 | 0.00078 |
| IAA | salt | 80 | 1.7 | 0.2 | 0.0 | 0.99 |
| IAA | dose | 80 | 6.2 | 0.015 | 10.1 | 0.0022 |
| IAA | salt $\times$ dose | 80 | 0.0 | 0.89 | 1.1 | 0.29 |

Alex Cantó-Pastor, Kaisa Kajala, Lidor Shaar-Moshe, Concepción Manzano, Prakash Timilsena, Damien De Bellis, Sharon Gray, Julia Holbein, He Yang, Sana Mohammad, Niba Nirmal, Kiran Suresh, Robertas Ursache, G. Alex Mason, Mona Gouran, Donnelly A. West, Alexander T.

*Figure placeholder: upload* `Figs/FigS_ISA3_celltype.pdf` *to this project.*

Figure S8: ***ISA3* transcripts are enriched in the columella/root-cap cluster of the *Arabidopsis* root, more specifically than other starch-metabolic genes.** (a) Proportion of cells with at least one *ISA3* UMI in each annotated cell type of GSE158761 (Serrano-Ron et al., 2021); dot area is the percentage of cells of that type, and cell types are ordered by size with the columella/root-cap cluster shaded. Genes are grouped as the candidate gene, other starch-metabolic genes, columella markers used as positive controls, and markers of other root cell types used as negative controls. (b) Fold enrichment in the columella/root-cap cluster, defined as the proportion of cells of that type with at least one UMI divided by the proportion across all 6,658 cells, in the same gene order; the dashed line marks no enrichment and $P$ values are from one-sided Fisher exact tests. The negative controls are strongly depleted in the columella/root-cap cluster, which excludes greater sequencing depth there as an explanation for the enrichment of *ISA3* and of the positive controls. Underlying values are given in Table S4.

Table S4: ***ISA3* is enriched in the columella/root-cap cluster of the *Arabidopsis* root single-cell atlas.** Cell-type enrichment in GSE158761 (Serrano-Ron et al., 2021), using the authors' own cell-type annotation (6,658 cells; 1,015 annotated as columella/root cap). For each gene we counted cells with at least one UMI, and compared the proportion detected within the columella/root-cap cluster with the proportion across all cells by a one-sided Fisher exact test. Fold is the ratio of those two proportions. Positive controls (*SMB*, *BRN2*) confirm the annotation; the negative controls are depleted in the same cluster, which rules out greater sequencing depth there as an explanation for enrichment. The equivalent test is not possible in tomato, where *ISA3* is detected at approximately one transcript per positive cell in root-tip single-cell data (Cantó-Pastor et al., 2024).

| Gene | Role | Cells detected | % of cluster | % of all cells | Fold | $P$ |
|---|---|---|---|---|---|---|
| *SMB* | root cap, positive control | 226 | 22.27 | 3.63 | 6.13 | $9.3 \times 10^{-172}$ |
| *BRN2* | distal columella, positive control | 157 | 15.47 | 2.51 | 6.17 | $9.1 \times 10^{-119}$ |
| *ISA3* | candidate gene | 62 | 6.11 | 1.82 | 3.36 | $9.5 \times 10^{-21}$ |
| *DPE2* | starch metabolism | 134 | 13.20 | 6.38 | 2.07 | $1.6 \times 10^{-18}$ |
| *AMY3* | starch metabolism | 52 | 5.12 | 2.49 | 2.05 | $1.0 \times 10^{-7}$ |
| *DPE1* | starch metabolism | 31 | 3.05 | 1.28 | 2.39 | $1.1 \times 10^{-6}$ |
| *SEX4* | starch metabolism | 15 | 1.48 | 0.50 | 2.98 | $3.5 \times 10^{-5}$ |
| *SGR9* | gravitropism | 2 | 0.20 | 0.77 | 0.26 | 1.00 |
| *PIN3* | negative control | 19 | 1.87 | 12.26 | 0.15 | 1.00 |
| *PIN7* | negative control | 10 | 0.99 | 5.50 | 0.18 | 1.00 |
| *SHR* | negative control | 2 | 0.20 | 3.05 | 0.06 | 1.00 |
| *LBD16* | lateral-root primordium, negative control | 5 | 0.49 | 14.76 | 0.03 | 1.00 |
| *WOX5* | quiescent centre, negative control | 0 | 0.00 | 1.37 | 0.00 | 1.00 |
| *GATA23* | lateral-root founder, negative control | 0 | 0.00 | 0.87 | 0.00 | 1.00 |
| *LBD29* | lateral-root primordium, negative control | 0 | 0.00 | 1.02 | 0.00 | 1.00 |

Borowsky, Kenneth A. Shackel, Neelima Sinha, Julia Bailey-Serres, Niko Geldner, Song Li, Rochus Benni Franke, and Siobhan M. Brady. A suberized exodermis is required for tomato drought tolerance. *Nature Plants*, 10(1):118–130, 2024. doi: 10.1038/s41477-023-01567-x.

Arjun Chandrasekhar and Magdalena M Julkowska. A mathematical framework for analyzing wild tomato root architecture. *Journal of Computational Biology*, 29(4):306–316, 2022.

Arjun Chandrasekhar and Saket Navlakha. Neural arbors are pareto optimal. *Proceedings of the Royal Society B*, 286(1902):20182727, 2019.

Wayne A. Fuller. *Measurement Error Models*. John Wiley & Sons, 1987.

Maryam Rahmati Ishka, Eric Craft, Miguel Pineros, and Magdalena M. Julkowska. Exogenous hormone treatments reveal species-specific regulation of individual components of root architecture and salt ion accumulation in cultivated and wild tomatoes. *Plant Direct*, 2026. doi: 10.1002/pld3.70169.

Laura Serrano-Ron, Pablo Perez-Garcia, Alvaro Sanchez-Corrionero, Inmaculada Gude, Javier Cabrera, Pui-Leng Ip, Kenneth D. Birnbaum, and Miguel A. Moreno-Risueno. Reconstruction of lateral root formation through single-cell rna sequencing reveals order of tissue initiation. *Molecular Plant*, 14(8):1362–1378, 2021. doi: 10.1016/j.molp.2021.05.028.

Pauli Virtanen, Ralf Gommers, Travis E. Oliphant, Matt Haberland, Tyler Reddy, David Cournapeau, Evgeni Burovski, Pearu Peterson, Warren Weckesser, Jonathan Bright, Stéfan J. van der Walt, Matthew Brett, Joshua Wilson, K. Jarrod Millman, Nikolay Mayorov, Andrew R. J. Nelson, Eric Jones, Robert Kern, Eric Larson, C J Carey, İlhan Polat, Yu Feng, Eric W. Moore, Jake VanderPlas, Denis Laxalde, Josef Perktold, Robert Cimrman, Ian Henriksen, E. A. Quintero, Charles R. Harris, Anne M. Archibald, Antônio H. Ribeiro, Fabian Pedregosa, Paul van Mulbregt, and SciPy 1.0 Contributors. SciPy 1.0: Fundamental Algorithms for Scientific Computing in Python. *Nature Methods*, 17:261–272, 2020. doi: 10.1038/s41592-019-0686-2.
